# Dynamic Reorganization of the Frontal Parietal Network During Cognitive Control and Episodic Memory

**DOI:** 10.1101/709220

**Authors:** Kimberly L Ray, J Daniel Ragland, Angus W MacDonald, James M Gold, Steven M Silverstein, Deanna M Barch, Cameron S Carter

**Affiliations:** Imaging Research Center, UC Davis, Sacramento, CA; Department of Psychology, University of Minnesota, Minneapolis, MN; Maryland Psychiatric Research Center, University of Maryland School of Medicine, Baltimore, MD; Department of Psychiatry, Rutgers - Robert Wood Johnson Medical School, Piscataway, NJ; Department of Psychiatry, Washington University, St Louis, MO; Department of Psychology, UC Davis, Davis, CA

**Author notes:** Corresponding Author: Cameron S Carter, 4701 X Street, Sacramento CA 95817.

## Abstract

Higher cognitive functioning is supported by adaptive reconfiguration of large-scale functional brain networks. Cognitive control (CC), which plays a vital role in flexibly guiding cognition and behavior in accordance with our goals, supports a range of executive functions via distributed brain networks. These networks process information dynamically and can be represented as functional connectivity changes between network elements.

Using graph theory, we explored context-dependent network reorganization in 56 healthy adults performing fMRI tasks from two cognitive domains that varied in CC and episodic-memory demands. We examined whole-brain modular structure during the DPX task, which engages proactive CC in the frontal-parietal cognitive control network (FPN), and the RiSE task, which manipulates CC demands at encoding and retrieval during episodic-memory processing, and engages FPN, the medial-temporal lobe and other memory related networks in a context dependent manner.

Analyses revealed different levels of network integration and segregation. High CC conditions exhibited greater integration across tasks. Moreover, nodes associated with higher cognitive functioning displayed the greatest amount of *dynamic* module reorganization across tasks. Specifically, the FPN displayed a high level of segregation in the DPX task, where only demands for proactive control varied, and more complex network integration with default mode and salience networks in the RiSE task, where CC is differentially integrated during memory encoding and retrieval. These findings provide insight into how brain networks reorganize to support differing task contexts, suggesting that the FPN flexibly segregates during focused proactive control and integrates to support control in other domains such as episodic-memory.

## Introduction

Cognitive control (CC) plays a vital role in flexibly guiding cognition and behavior in accordance with our goals and it is thought that this ability serves as an important element in healthy brain function (Veen & Carter, 2006). This mechanism is not limited to a particular cognitive domain (Banich, 1997), CC supports a range of cognitive functions, including working memory, episodic memory (Ragland et al., 2009), inhibitory processing (Banich et al., 2000), and goal maintenance (Henderson et al., 2012). These aspects of executive functions are supported by distributed brain networks that represent and process information in a dynamic manner via functional connectivity (FC) between network elements (Cole & Schneider, 2007; Cole et al., 2013).

The prefrontal cortex (PFC) plays a central role in cognitive control (Badre, 2008; MacDonald, 2000; Niendam et al., 2012). Evidence supports an anterior-posterior gradient of function within the PFC. While the rostrolateral PFC is associated with relational reasoning, functional magnetic resonance imaging (fMRI) studies suggest that interactions between dorsal and ventral lateral prefrontal regions and posterior brain regions including the lateral parietal lobe and the medial temporal lobe (MTL), support the retrieval of relationally encoded information and associative recognition during episodic memory(Murray & Ranganath, 2007; Ragland et al., 2012; 2015). More broadly, the evidence for such segregated brain activity during the processing of goal maintenance is consistent with recent studies suggesting that trial-by-trial cognitive control engages a large-scale functional brain network encompassing frontal and parietal cortices (Henderson et al., 2012; Lopez-Garcia et al., 2015).

Advances in applying graph analysis to fMRI data have provided means to mathematically describe and quantify cognition related patterns of function connectivity. Initially restricted to rest, developments in graph theory (GT) methodology applied to task fMRI have furthered our understanding of how cognitive control is supported through flexible, context-dependent integration and segregation of functional brain networks in a dynamic manner (Braun et al., 2015; Cocchi et al., 2013a; Cocchi et al., 2013b; Fornito et al., 2012; Hearne et al., 2017). These studies commonly focus on network organization supporting performance of a single cognitive task within a single brain network or defined set of networks. More recently, studies have demonstrated that modular properties of brain networks shift in response to differing cognitive demands (Cohen et al., 2016; Geib et al., 2017, Westphal et al., 2017). The current fMRI study examines whole brain network organization involved in cognitive control during two tasks that involve multiple cognitive domains. We examine network reorganization specifically within the framework of changes in functional connectivity where we investigate two different mechanisms of reorganization: (1) enhanced network connectivity in the frontal parietal network (FPN) using the participation coefficient and (2) enhanced between network connectivity, using modularity, where cognitive systems integrate to create new networks.

We use modularity, a GT method that measures the decomposability of a graph into modules or communities, to provide insight into context-dependent network reorganization in healthy adults (HC) performing tasks with varying demands on cognitive control. Modularity is a general hallmark of complex biological systems. Modular organization of brain networks shapes how information is distributed and processed where regions that are functionally close and tend to share information are considered members of the same cluster or module (Sporns & Betzel, 2016). A module containing nodes from a variety of cognitive systems likely indicates functional integration amongst cognitive brain networks, whereas a module composed of only nodes from a single system may likely represent network segregation.

Reliability of the brain’s functional architecture at rest is well supported, however recent GT analyses have revealed an adaptive reconfiguration of large-scale brain networks that support higher cognitive functions (Bassett et al., 2011; Cole et al., 2014; Braun et al., 2015; Hearne et al., 2017). Studies probing the correspondence of network organization during rest and task using residualized and nonresidualized fMRI data have found that network modifications during task were independently associated with regional activation and changes in functional connectivity (Gratton, Laumann, Gordon, Adeyemo, & Petersen, 2016). Ultimately this suggests that meaningful, context-dependent network reconfigurations occur against a backdrop of stable, large-scale networks that support diverse cognitive functions (Cohen & D’Esposito, 2016; Cole et al., 2014; Crossley et al., 2013; Hearne et al., 2017).

The current study examines cognitive control processing in data from 56 healthy adults performing fMRI tasks from two distinct cognitive domains that varied in demands for cognitive control, the RiSE episodic memory task and the Dot Pattern Expectancy (DPX) goal maintenance task. Adapting a beta series correlation technique (Mumford et al., 2012) to examine brain wide integration and segregation of cognitive systems during the RiSE and DPX, we leverage opposing network topology to *quantitatively* assess the dynamic network reorganization (i.e. variation in connectivity states) involved in each of these cognitive tasks. Prior work from this sample used the Network Based Statistic (Zalesky et al., 2010) to identify increases in functional connections within the FPN associated with cognitive control demand (Ray et al., 2017). However, it is not clear how within network FC changes correspond to brain wide network organization. Here, we investigate context-dependent network reorganization via changes in integration and segregation properties associated with modular organization across the RiSE and DPX tasks. We predict that increased demands for cognitive control processing will result in greater brain wide integration of cognitive systems measured by decreased modularity. We measure changes in modular partitions associated with increased CC using mutual information, We hypothesize that the FPN will uniquely integrate with other higher-cognitive networks in order to support efficient cognitive processing/task completion which we quantify by means of increased participation coefficient of modular partitions. Finally, we expect that our hypothesized increased integration during the RiSE task will correspond with a merging of FPN and memory related systems.

## Materials and Methods

### Subjects

Study participants were recruited as part of the CNTRACS Consortium (http://cntracs.ucdavis.edu), which included 5 different research sites: University of California—Davis, Maryland Psychiatric Research Center at the University of Maryland, Rutgers University – Robert Wood Johnson Medical School, University of Minnesota—Twin Cities, and Washington University. Recruitment and informed consent procedures for each site were approved by their Institutional Review Boards. Complete details regarding CNTRACS recruitment and enrollment can be found in Ragland et al. (2015).

Data were obtained on 60 healthy adults (HC). Participants were excluded if they exhibited excess movement (i.e., >0.37 mm mean frame-to-frame movement), below-chance performance, or image acquisition errors. This left final samples of 56 HC (34.0±11.4 yrs) for the RiSE task, and 52 HC (34.1±10.4 yrs) for DPX task (Table 1).

**Table 1:**
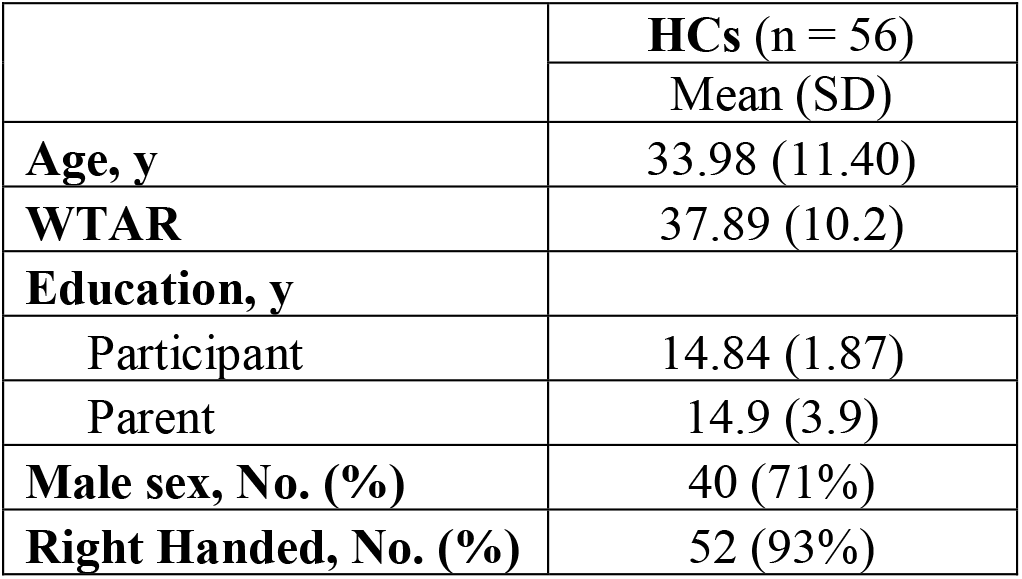
Participant Demographics. Abbreviations: HCs, healthy controls; WTAR, Wechsler Test of Adult Reading.

While fMRI and behavioral data from these subjects have been used in previous publications, results from the current study are unique and do not include previously published findings.

### Data Acquisition

#### Relational and Item-Specific Encoding (RiSE) task

The design was identical to that of the original RiSE studies (Ragland et al., 2012; 2015), with the following exceptions: stimuli were presented in pairs during both encoding conditions (see below), and the item recognition task did not include confidence ratings. Participants completed 4 encoding and 4 retrieval fMRI runs. During encoding (Figure 1A), participants alternated between 3 item-specific blocks (“Is either object living?”; 9 low cognitive control trials each) and 3 relational blocks (“Can one object fit inside the other?”; 9 high cognitive control trials each) in a “jittered” event-related design. During item recognition (Figure 1B), participants made a 2-button response to indicate whether objects were previously studied (old) or never studied (new). During item recognition, 54 individual objects from each encoding condition (54 item-specific, 54 relational) were randomly presented with 54 new items. The Rise task is considered a of rapid-presentation event-related fMRI paradigm, therefore OPTSEQ (available at https://surfer.nmr.mgh.harvard.edu/optseq/) was used to optimize the efficiency of trial presentation timing and randomization across each block. Because our interest was in engagement of CC during encoding processes rather than accuracy of frequently equivocal responses (eg, Is an apple that is not on the tree living?), fMRI analysis included trials in which participants correctly responded to during the recognition condition and their corresponding encoding trials. As reported in Ragland et al., 2015, mean accuracy for healthy adults was 72.0% and 86.1% for item and relational recognition trials respectively. See (Ragland et al., 2015) for more information regarding the RiSE task.

**Figure 1:**
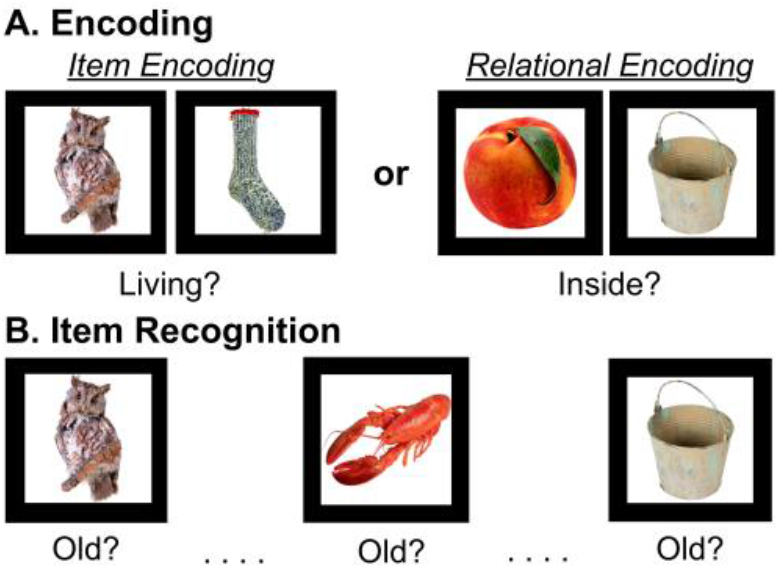
Illustration of the RiSE task. A) Item specific and relational object pairs presented while subjects made either an item-specific encoding response or relational encoding response. B) During item recognition, objects from item and relational encoding conditions were randomly presented with new items, and participants indicated whether each item was old (i.e. previously studied).

Previous RiSE fMRI studies contrasting relational (high cognitive control) against item-specific (low cognitive control) in the same sample have identified robust activation increases in the bilateral DLPFC, VLPFC, parietal, and occipital cortices (Ragland et al., 2015). Furthermore, functional connectivity analyses examined in this sample have demonstrated network specific engagement of the FPN during the RiSE task (Ray et al., 2017).

#### Dot Pattern Expectancy (DPX) task

The DPX task consisted of a sequence of cue-probe stimuli where participants made one response when a target cue-probe pair was presented and another response for all other stimuli (Figure 2). Cues indicated the need for high (B Cues) or low (A Cues) levels of cognitive control. Four types of trials were presented across four blocks: AX, AY, BX and BY. AX trials are “target trials” where a valid cue is followed by a valid probe. The 3 other trial types are “Non-target trials” in which either a valid cue is followed by an invalid probe (“AY” trials) or an invalid cue is followed by either a valid or invalid probe (“BX” or “BY” probes, respectively). Each block of the DPX task consisted of 40 trials: 24 AX, 6 AY, 6 BX, and 4 BY. The nature of the cue (valid or invalid) provides the “context” for responding on a given trial. The majority of trials are “target trials” (AX trials). This feature is intended to encourage participants to “expect” a valid probe to follow a valid cue. A consequence of this manipulation is that participants develop a prepotency to respond with “target” responses on trials for which valid cues are presented, regardless of whether the trials were of the target (AX) or non-target (AY) type. The DPX task is a rapid-presentation event-related fMRI paradigm, therefore OPTSEQ (available at https://surfer.nmr.mgh.harvard.edu/optseq/) was used to optimize the efficiency of trial presentation timing and randomization across each block. Correct responses from 4 runs of the DPX task that used for analysis. See (Poppe et al., 2016) for more detail regarding the AX-DPX.

**Figure 2:**
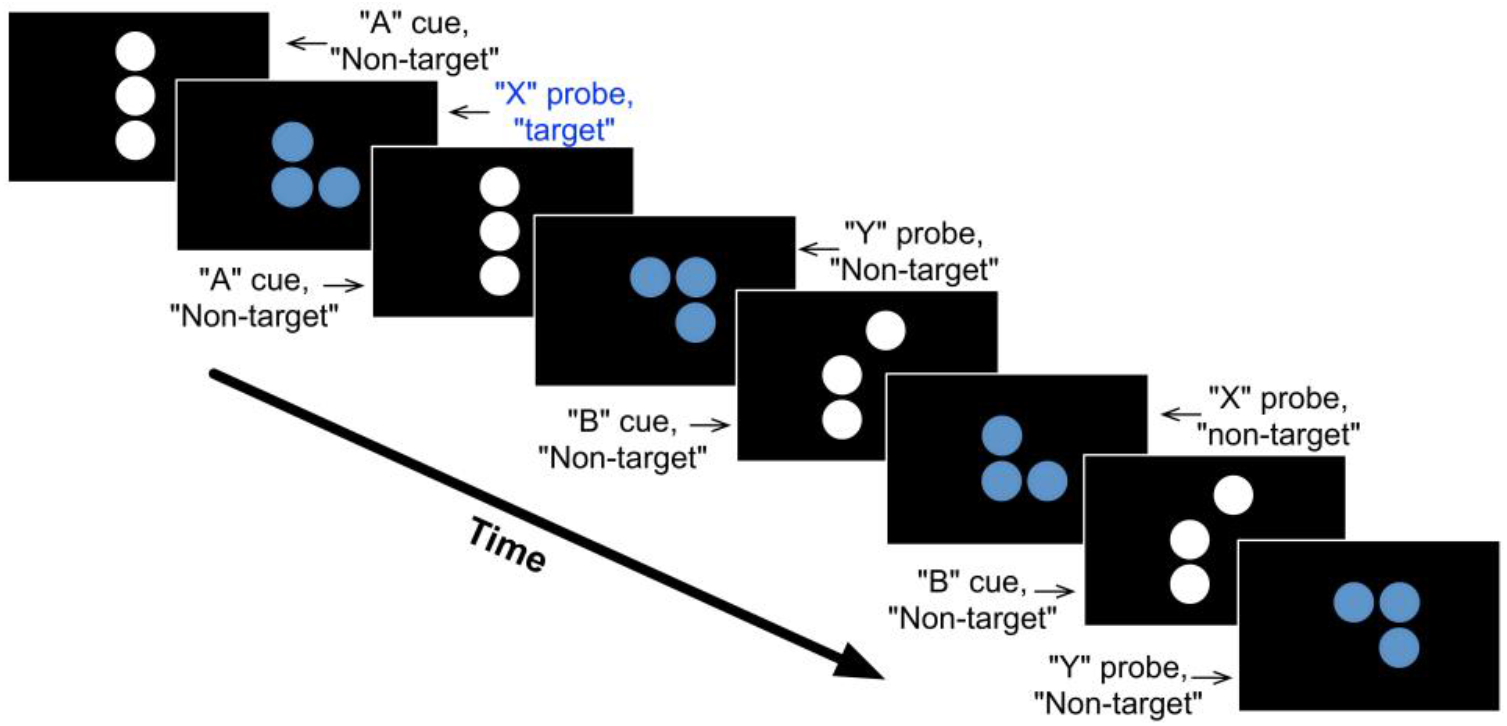
Illustration of the Dot Pattern Expectancy Task. Shown is an example sequence of cue-probe stimuli and the type of response (target or non-target) a participant was required to make after each stimulus. The nomenclature for stimuli and trial types was adopted from the expectancy letter AX task. The valid cue pattern is referred to as ‘‘A’’ and the valid probe pattern is referred to as ‘‘X’’. Non- “A” cue patterns are referred to as “B”-type cues, and non-“X” probe patterns are referred to as “Y”-type probes. A target response is required to “X” when it follows “A”, non-target responses are made after all other stimuli. The first pair of stimuli in the sequence represents an AX trial. The third and fourth stimuli together represent an AY type of trial, the fifth and sixth stimuli together complete a BX trial, and the seventh and eighth stimuli make up a BY type of trial.

Previous fMRI whole-brain analyses from our group using a different healthy sample have shown that contrasting B-cues relative to A-cues in the DPX task elicits widespread activations in the cognitive control FPN, including bilateral DLPFC, bilateral fusiform gyri, and right inferior parietal gyrus (Lopez-Garcia et al., 2015). These activation findings have been replicated in the current sample (Poppe et al., 2016) and subsequent functional connectivity analyses have also demonstrated network specific engagement of the FPN during the DPX task (Ray et al., 2017).

### Preprocessing

Images were acquired in a single 3T MRI session using a consistent protocol across sites. Functional images were acquired using gradient-echo BOLD echo-planar imaging (TR=2000 ms, TE=30 ms, 77° flip angle, FOV=220 mm^2^, 3.43×3.43×4 mm voxels, 32 axial slices parallel with the anterior/posterior commissure). For more information see (Henderson et al., 2012).

Pre-processing was carried out using the FMRI Expert Analysis Tool (FEAT) in the FMRIB Software Library (FSL version 4.1; www.fmrib.ox.ac.uk/fsl) using standard procedures, including fieldmap correction, spatial normalization and nonlinear registration to MNI152. Field maps to correct fMRI data for geometric distortion caused by magnetic field inhomogeneities and a T1-weighted anatomical image (1-mm isotropic voxels) were also acquired.

### Data Processing (Beta-series regression)

Subject-wise beta-series regression analysis was performed on RiSE and DPX fMRI data in order to capture trial specific BOLD effects for each condition (Turner et al., 2012). Using the least squares-separate (LS-S) method to measure event-related functional connectivity, individual trials were modeled with a new GLM in SPM8 with two predicted BOLD timecourses—one that reflects the expected BOLD response to the current event and another for the BOLD responses to all events except the current event. All events were modeled, but only cue events for DPX correct trials and correct RiSE trials were included in the analysis, because these represented trial periods in which cognitive control demands were maximized.

Separate regressors modeling each event were defined in a general linear model to yield unique condition-wise beta values for every voxel. Each beta value reflected the magnitude of the hemodynamic response evoked by each event. Beta images were sorted by condition and concatenated across runs yielding a 4D dataset (space x n trials), or *beta-series*, for each of the six conditions: RiSE *Item* Encoding, RiSE *Relation* Encoding, RiSE *Item* Recognition, RiSE *Relation* Recognition, DPX A Cue, and DPX B Cue.

Additional motion correction steps were taken analogous to data-scrubbing procedures often performed in resting-state functional connectivity analyses (Power et al., 2014). Beta-images containing frame-wise displacement (FD) values greater than 0.5mm motion were excluded from beta-series. If more than 10% of TRs within a block contained frame-wise displacement >0.5 mm, the entire block was excluded from analysis. This FD threshold led to the exclusion of 5 blocks of the RiSE task, and 1 block in the DPX task.

Next, each participant’s brain data was parcellated into discrete regions of interest representing nodes obtained from the Power atlas (Power et al., 2011). Twenty-one Power nodes were eliminated due to low signal, and two bilateral MTL nodes were added (MNI coords: −30,−12,−22; 32,−14,−22) resulting with 245 nodes across the whole-brain. Beta-series pairwise correlations for all 245 nodes were extracted and z-transformed resulting with a 245 by 245 connectivity matrix. A final group FC graph for each condition was established by summing the 5% thresholded connection matrices across all subjects, then applying a subsequent 5% threshold on the summed group connection matrix to identify connections that are consistently strongest across participants, followed by binarization of the summed group connection matrix.

### Graph Analysis

Recent advances in the application of graph theoretical analysis to fMRI data allow us to leverage information contained within the BOLD signal to test hypotheses regarding the functional architecture of the human brain (Bullmore & Bassett, 2011). One well-known investigation of community structure in functional brain networks (Power et al. 2011) extracted communities from resting-state fMRI data and, using a map of task-based activations across a range of tasks, mapped these communities to well-studied cognitive systems. The current study examines segregation and integration of brain networks via modular organization during each condition in the RiSE and DPX tasks separately and leverages opposing network topology to highlight the dynamic reorganizations that support these different cognitive control tasks in healthy adults. That is, we used graph theoretical measures (i.e. modularity) to model and quantify context dependent changes in functional brain organization during three fast event-related fMRI paradigms: The RiSE encoding, RiSE recognition, and DPX tasks.

#### Modularity

Modularity (Q) and modular partitions were extracted for each subject using the community Louvain algorithm provided by the Brain Connectivity toolbox (Rubinov & Sporns, 2010). A Louvain module partition yields a set of non-overlapping communities, in which each network node belongs to one and only one module. Modularity ranges from zero to one and quantifies the goodness of modular partitions, where good modularity partitions have high modularity values and an unexpectedly high proportion of connections within modules, and an unexpectedly low proportion of connections between modules (Newman, 2004). A high proportion of connections between modules (i.e. low modularity) suggests a greater level of brain wide community integration amongst cognitive systems, whereas a low proportion of connections between modules (i.e. high modularity) may suggest a segregation of communities. Nevertheless, the Louvain modularity algorithm uses a randomized heuristic approach and consequently results across runs slightly vary. We therefore applied this algorithm 1,000 times for each task condition at proportional thresholds of 0.05 through 0.60 (to avoid negative connections in FC matrices) at increments of 0.05, and selected the partition with the highest modularity score, Q (Rubinov & Sporns, 2011; Meunier, 2009). Modularity analyses were run on the group connection matrix and on individual subject connection matrices for each task.

#### Comparison of task-based Q to null model

Null models are important adjuncts of descriptive graph analysis as they allow discriminating which graph attributes are due to chance, and which exceed the expected values given by the null model (Sporns, 2018). To establish a null-model for each condition examined, we assessed the modularity of random graphs derived from each task condition. Random graphs for each participant and for the group mean were created using functions provided in the Brain Connectivity Toolbox that randomize connections in their corresponding task-based network (randmio_und.m, 500 iterations), while preserving the degree distribution (Rubinov & Sporns, 2011). Modularity (Q) of task-based graphs were compared to their respective null-model across a range of thresholds (i.e. 0.05 – 0.60 proportional thresholds at intervals of 0.05) using a repeated measures ANOVA.

#### Comparison of task-based Q to Power Q

An advantage of using the Power atlas is that it also provides an *a priori* partition of subgraphs that replicate across cohorts and correspond anatomically with many functional systems consistently observed in the neuroimaging literature (Power et al., 2011). As a follow up to task-based modularity analyses of the RiSE and DPX paradigms, we were interested in comparing the task-based Louvain-Modular partitions to Power et al.’s, resting-state subgraph partition. To do so, modularity (Q) of the Power resting-state subgraph partition was calculated for each subject using an abridged version of modularity scripts provided in the BCT that determines Q based on the input Power partition rather than an optimized partition. Modularity of the Power partition for each subject was then compared to the modularity of their corresponding task-based partitions using a repeated measures ANOVA.

#### Quantifying Network Reorganization using Mutual Information Theory

Finally, we utilized a popular information-theoretic measure of distance in partition space, mutual information (Meilă, 2007; Rubinov & Sporns, 2011), to provide a measure of the amount of network reorganization between high and low control condition partitions in the RiSE and DPX tasks. As described by Rubinov and Sporns (2011), mutual information between two partitions *M* and *M*’ was calculated as:

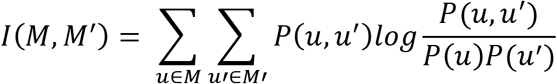

where 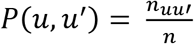 and *n*_*uu*′_, is the number of nodes that are simultaneously in module *u* of partition *M*, and in module *u*’ of partition *M*’. Partition vectors *M* and *M*’ contained the full set of 245 nodes, thus mutual information values reflect network reorganization across the whole-brain. In this context of the current study, mutual information indicates the degree to which nodes are similarly assigned to modules during high and low cognitive control conditions. A mutual information value of one would indicate two identical partitions are being compared and no change in module organization between high and low cognitive control conditions. A mutual information index of zero would indicate maximally different partitions and thus a large change in module organizations between the two conditions compared.

#### Participation Coefficient

We utilized the participation coefficient (PC) to examine the integrative role of the FPN with respect to other cognitive networks. The participation coefficient measures the proportion of connections a node has within its own module versus other modules in the network (Guimerà et al., 2005; Rubinov & Sporns, 2010; Sporns et al., 2007). Thus, nodes with high PC are more strongly connected to nodes associated with other modules, thereby facilitating greater integration of information across modules; in contrast, nodes with lower PC are predominately connected to nodes within its assigned module. The PCs of FPN nodes in the Power atlas were extracted from the 5% thresholded functional connectivity matrices of each task condition for each individual participant. PCs were averaged across all nodes within the FPN, and a repeated measures 3×2 ANOVA was performed to test the effect of cognitive control across the RiSE encoding, RiSE recognition, and DPX tasks.

## Results

### Comparison of task-based Q to null model

Subject-wise functional partitions thresholded to the strongest 5% of connections were significantly more modular (Q) than their respective null models for each task, *F*(1,51)=2497,*p*<0.001(Fig 3). This significant difference was present across a range of thresholds.

**Figure 3:**
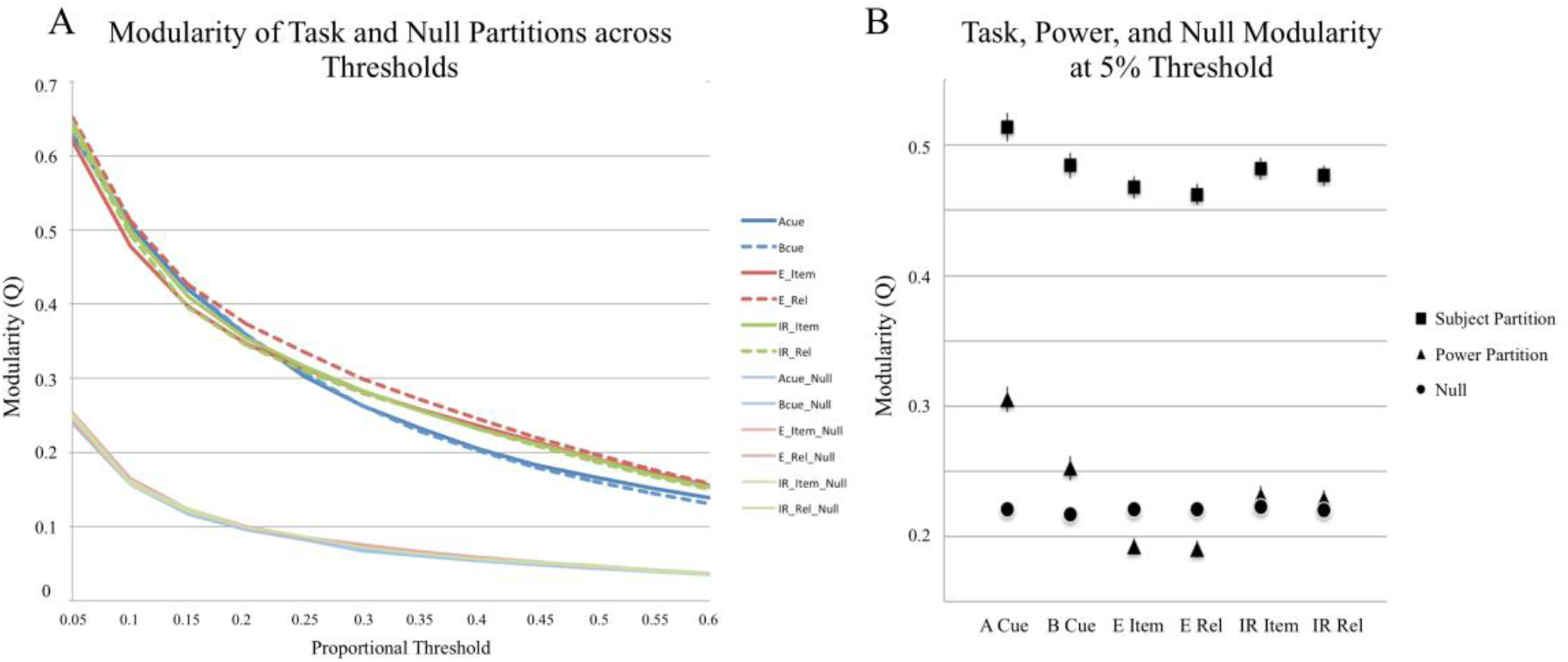
A) Louvain modularity(Q) of group mean task-based Louvain partitions compared to their respective individual null-model. Task-based partitions were significantly more modular than their null model across all thresholds tested (*p*<0.001). B) Modularity of each individual’s task-based partitions at a 0.05 proportional threshold compared to their null-model and the Power subgraph partition.

### Comparison of task-based Q to Power Q

Compared to modularity (Q) of the *a priori* Power partition, we found that the task-based functional partitions provided a significantly greater modularity score than the resting-state subgraph partition (F(1,51)=9710.173, *p*<0.001; 5% threshold). A greater Q-score indicates that newly established task modules yield a better model of network organization during the RiSE and DPX paradigms than the resting-state Power subgraph partition, and that a reasonable amount of network reconfiguration occurs between rest and task (Hearne et al., 2017). Moreover, Power modularity scores (Q) were, on average, 0.013 higher than their null models across tasks (F(1,51)=6.541, *p*=0.014; 5% threshold).

### Task-based Modularity

Once the non-randomness of task-based modular partitions was established across participants, we performed a 3×2 ANOVA to examine effects of task and cognitive control. In doing so, we observed an overall effect of task (F(2,50) = 9.615, p < 0.001), indicating that the RiSE and DPX tasks exhibit different levels of network integration and segregation (Fig 3B). A cognitive control effect was also present, where low cognitive control conditions exhibited greater modularity relative to high control conditions across RiSE and DPX conditions (F(1, 51)=5.673, p < 0.021). Furthermore, a task by control interaction effect was present (F(1,51)=3.179, p < 0.046), indicating that while low cognitive control conditions were greater than high cognitive control conditions in each of the tasks examined, that the extent of this modularity effect varied across tasks.

### Quantifying Network Reorganization using Information Theory

While modularity can be used as an estimate of whole brain integration or segregation amongst brain modules, it cannot capture module composition or changes in modular partitions. Thus, *we used mutual information to quantify the similarity of module assignments between high and low cognitive control conditions* in the RiSE encoding, RiSE recognition, and DPX tasks, which we interpret as a measure of network reorganization.

A repeated measures ANOVA examining the mutual information scores comparing high and low cognitive control condition partitions in the RiSE encoding, RiSE recognition, and DPX tasks identified a significant effect of task (F(2,51) = 94.558, p <0.001; Fig 4), indicating that different levels of network reorganization were observed across tasks. Post-hoc t-tests of mutual information scores identified significant differences between each task: DPX > RiSE_Recognition_ (t = 5.726, p > 0.001), DPX > RiSE_Encoding_ (t = 12.436, p > 0.001), RiSE_Recognition_ > RiSE_Encoding_ (t = 8.673, p <0.001). It can be seen in Figure 4 that the greatest agreement between modular partitions for high and low control conditions (quantified via Mutual Information) was observed in the DPX task, followed by RiSE recognition, with the least agreement found for RiSE encoding. Considering that larger mutual information scores indicate fewer differences between two modular partitions, we infer that the reconfiguration of brain networks was lowest between the DPX conditions and greatest during the RiSE encoding conditions.

**Figure 4:**
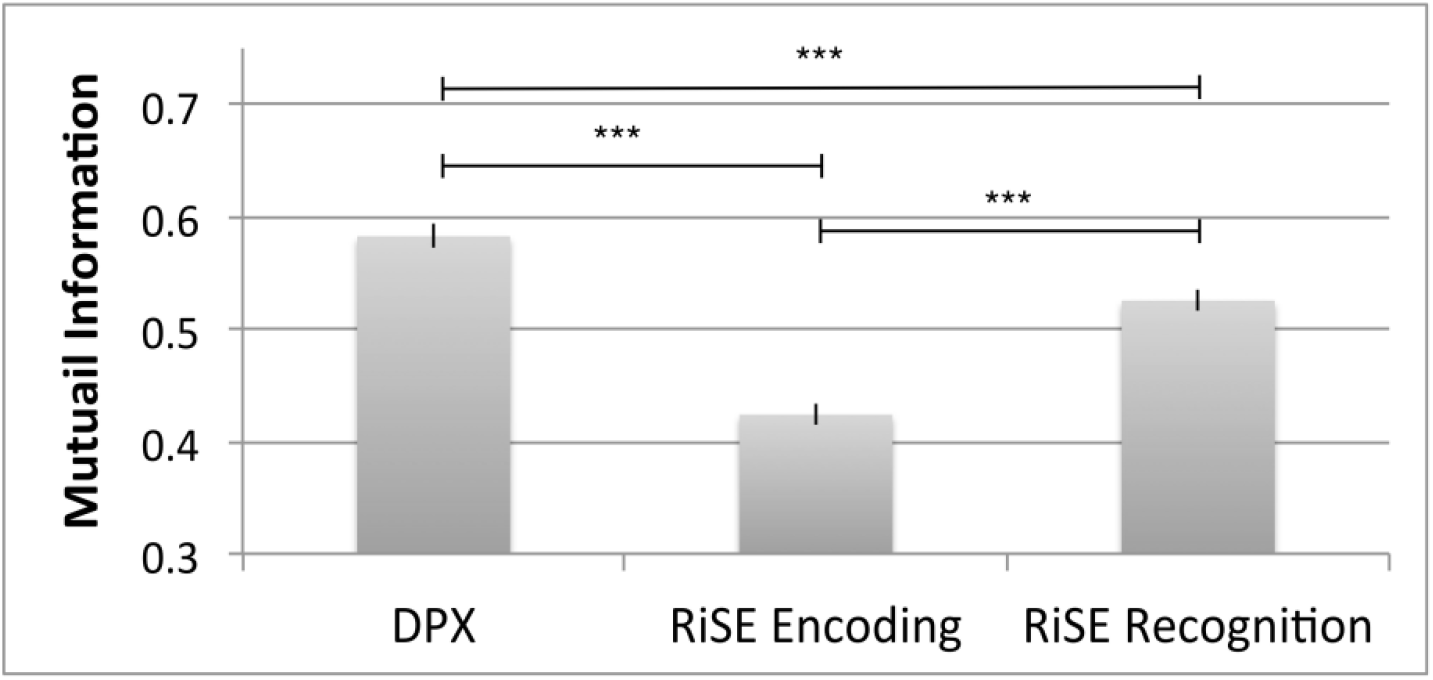
Mutual information scores quantifying the similarity of module partitions between high and low cognitive control conditions in the DPX, RiSE encoding, and RiSE recognition tasks. *** indicates p < 0.001.

### Participation Coefficient

We used the participation coefficient to assess changes in the diversity of intermodular connections within the FPN associated with cognitive control. We observed a main effect of cognitive control across the 3 tasks examined where PC was greater during high cognitive control conditions compared to low cognitive control conditions (F(2,51) = 4.083, p = 0.046). This finding indicates that the FPN exhibits greater between module communication (i.e. integration) during high cognitive control conditions relative to low cognitive control conditions.

#### INTEGRATION AND SEGREGATION OF COGNITIVE SYSTEMS

Varying levels of integration and segregation between cognitive systems were observed across the tasks examined. Visual inspection of partitioned brain graphs (Figure 5) revealed that nodes associated with low-level perceptual functioning were consistently constrained to two similar modules across tasks. This included nodes associated with sensorimotor (hand-royal blue, mouth-navy), auditory (orange), and cerebellar networks (yellow). Nodes associated with the visual system (red) displayed moderate levels of network reorganization, where medial nodes flexibly integrated with either lateral vision nodes during the DPX task, or posterior DMN (gray) and memory (cyan) nodes in the RiSE encoding task (Figure 6). Nodes associated with high-level cognitive functioning displayed complex levels of module reorganization across tasks. Notably, the FPN (spring green) displayed varying levels of integration and segregation across tasks as it uniquely integrated with DMN and salience (sky blue) networks in multiple modules across the RiSE conditions, but segregated into the main element of a single module in both DPX conditions.

**Figure 5:**
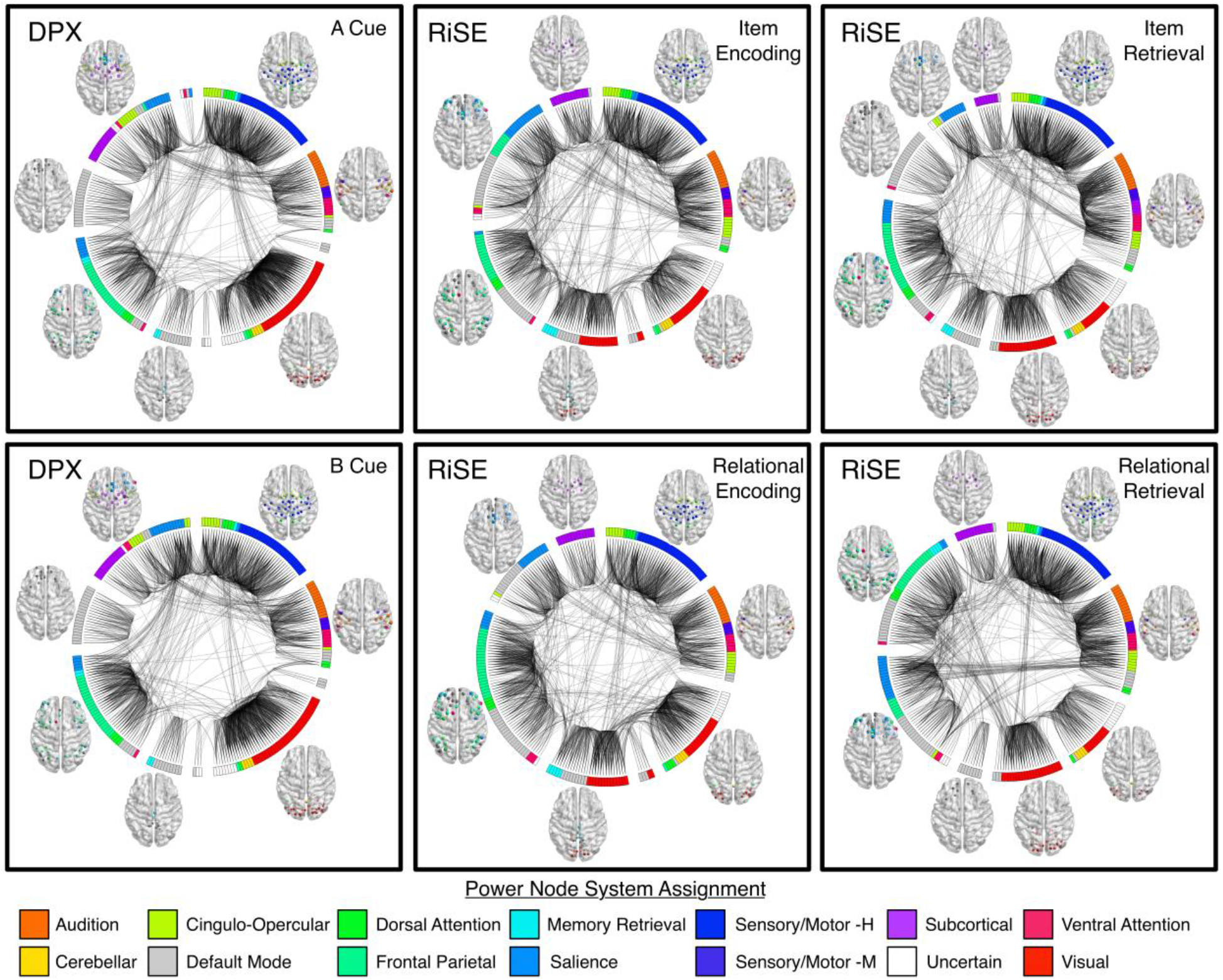
Brain graphs of healthy adults performing the DPX (left column), RiSE encoding (middle column), and RiSE recognition (right column) tasks. Low cognitive control conditions for each task are shown on the top row, high cognitive control conditions are shown on the bottom row. Edges displayed in each graph represent the strongest 5% of functional connections, groups of nodes indicate their module assignment defined using the Louvain Modularity algorithm in the BCT. The color of each node corresponds to one of the 14 cognitive systems proposed in Power et al., (2011). The layout of graphs were visualized using BrainNet and Circos.

**Figure 6:**
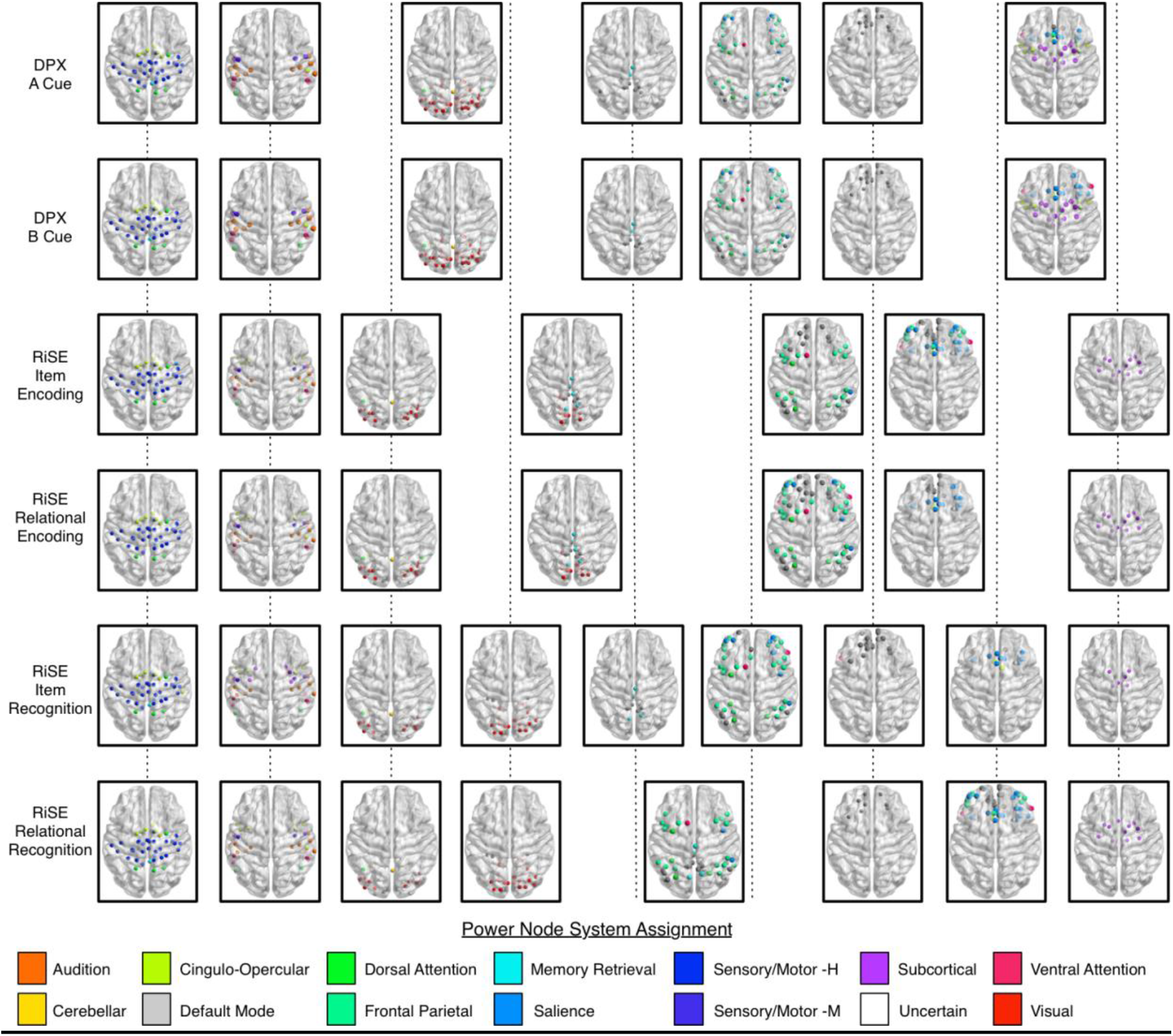
Brain modules of healthy adults performing the DPX, RiSE encoding, and RiSE recognition tasks. Each row represents the modular partition (i.e. the collection of modules in a brain graph) of a single condition during DPX (rows 1 and 2) and RiSE tasks (rows 3-6). Modules that align vertically across multiple rows (i.e. in the same column or along a common vertical dotted line) indicate that they are similarly organized across multiple task conditions. Modules that partially overlap vertically indicate an integration or segregation of cognitive systems into newly established modules across tasks. Node color scheme corresponds to cognitive systems proposed in Power et al., (2011). Brain Images were created using the BrainNet toolbox.

## Discussion

The purpose of the current study was to quantify and characterize changes in whole-brain network dynamics as they pertain to cognitive control. While previous functional connectivity studies have largely focused on changes within a single cognitive system, or between a set of systems (Dosenbach et al., 2008; Fornito et al., 2012; Hearne et al., 2015) in the present study we explored context-dependent integration and segregation of brain systems in healthy adults performing fMRI tasks from two distinct cognitive domains that varied in demands for cognitive control as well as demands for episodic mnemonic functions. In doing so, we leverage opposing community structure in the RiSE and DPX tasks to highlight the dynamic reorganizations that support these differentially engaging cognitive control tasks.

### Task based modularity

Examining modularity of the RiSE and DPX tasks revealed different levels of network integration and segregation. Across all tasks, modularity was lower in high control relative to low control conditions. Decreased modularity indicates a reduced proportion of within module connections in a network, and thus a greater proportion of between module connections suggesting a greater level of brain wide community integration. Similar findings of increased integration have been reported in a range of executive functions including inhibition (Spielberg et al., 2015) and deductive reasoning (Cocchi et al., 2013a; Hearne et al., 2017), and in episodic memory where lower whole-brain modularity has been associated with improved performance (Cohen & D’Esposito, 2016; Geib et al., 2017; Westphal et al., 2017). Interestingly, a task by control interaction effect was also present, indicating that while modularity was greater in low cognitive control conditions, the extent of the effect varied across tasks. Interaction effects suggest that modules were more segregated in the DPX task than in the RiSE recognition and encoding tasks. This finding is not surprising considering that the DPX primarily engages cognitive control, where as the RiSE task requires recruitment of both control and long-term (episodic) memory systems. Cohen and colleagues report similar reduced modularity in a memory task, which is thought to require coordination across multiple brain networks, compared to a motor task which likely involves only a single brain network (Cohen & D’Esposito, 2016).

We also found that the task-based functional partitions provided a significantly greater modularity score than the resting-state Power partition. Most comparisons between task and rest FC have observed high correspondence (Cole et al., 2014; Fair et al., 2007; Fox & Raichle, 2007; Greicius et al., 2008). However, more recent comparisons between task and rest functional connectivity have emphasized differences in connectivity patterns (Buckner et al., 2013; Hermundstad et al., 2013; Mennes et al., 2012). Although our sample does not include resting-state fMRI, it is clear that the task-based community structure we observed does not fit the task-negative partition proposed by Power and colleagues (2011). This is further supported by a recent study from Hearne and colleagues showing that increases in reasoning complexity resulted in a merging of resting state modules (Hearne et al., 2017). One source of this disagreement between rest and task community partitions may be due to differences in methods for community detection (Infomap compared to Louvain modularity), furthermore the beta-series approach used in the current analysis does not regress out task structure. Nevertheless the disparity between the presented task based modular partitions and the proposed Power resting-state subgraph partition do provide support for the concept of differences between task and resting functional architectures.

### Network Reorganization

Different levels of network reorganization were observed across tasks. Using mutual information as a measure of dynamic network reorganization between modular partitions of high and low cognitive control conditions, we observed the highest mutual information (i.e. agreement in module composition) in the DPX task, followed by RiSE recognition, and the least mutual information during the RiSE encoding. While there was no significant correlation between mutual information (I) between high and low CC conditions and modularity scores (Q) for any of the task conditions examined, similar effects across tasks were present. In other words, the DPX was most modular (Q) and contained the greatest mutual information (I) whereas RiSE encoding was least modular (Q) and contained the least amount of mutual information (I). Incorporating the current findings with previous studies investigating FPN specific functional connectivity changes associated with cognitive control on the same sample (Ray et al. (2017), the DPX task displayed the greatest change in within FPN functional connectivity between high and low CC conditions (Ray et al., 2017) and our current work suggests it is also most modular (i.e. greatest network segregation) and experiences the least amount of network reorganization. Conversely, the RiSE encoding task displayed the least within FPN change in function connectivity between high and low CC conditions (Ray et al., 2017) while current work shows RiSE encoding to be least modular yet undergoes the most network reorganization. All the while, performance on these tasks is significantly correlated, suggesting that they share common processes associated with cognitive control (Sheffield et al., 2014) and they elicit increased activity in the dorsolateral prefrontal and parietal cortices in fMRI studies(Lopez-Garcia et al., 2015; Ragland et al., 2015). Together, this may suggest that cognitive control processing may be recruited via mechanisms that enhance within network and between network functional connectivity.

### Integration and segregation of cognitive systems

The FPN exhibited greater integrative properties during high CC control conditions than low CC as measured by the participation coefficient. Subsequent visual inspection of group partitions was performed to better understand changes in module compositions corresponding changes in modularity (Q), mutual information (I), and participation coefficient (PC) across the DPX and RiSE tasks.

Across all conditions examined, several modules were consistently identified which included nodes associated with low-level perceptual systems (e.g. sensorimotor, cerebellar, audition, vision). Nodes in the FPN, DMN, and salience network varied in their module assignments. Similar findings have been reported in a whole-brain, task-based graph theory analysis comparing modular organizations between 2-back and 0-back conditions in a working memory, executive function n-back task (Braun et al., 2015). Braun and colleagues found that ‘flexibility’, a term defined as the tendency for nodes to change module allegiance, was highest in PFC-related systems, which corresponded with PFC module reorganization, observed in the DPX and RiSE tasks. In relation to episodic memory, Westphal and colleagues report an inverse relationship between DMN and FPN coupling (i.e. network integration) and whole-brain modularity during an episodic memory task (Westphal et al., 2017). This corresponds well with the RiSE task community structure where modules with FPN nodes also contain more DMN nodes and exhibit lower modularity relative to DPX community structure.

The notion of context-dependent dynamic integration and segregation of higher cognitive systems has recently been introduced by other investigative groups. For example Menon and colleagues have suggested that the activity of the frontal parietal and cingulo-opercular systems is related to the behavior of the DMN (Bressler & Menon, 2010; Menon & Uddin, 2010). The DMN is commonly referred to as ‘task negative’ because its activity decreases during external goal-oriented actions and increases during performance of tasks requiring self-related processing (Raichle & Snyder, 2007). Several early studies have supported the hypothesis of a functional antagonism between the two task-positive FPN and cingulo-opercular systems on the one hand and the task-negative DMN on the other (Kelly et al., 2008). Decreased activity within the DMN is inversely correlated with cognitive control (Lawrence et al., 2003; McKiernan et al., 2003) and positively correlated with task-unrelated mental activity (Mason et al., 2007) suggesting that they would be segregated systems. Recent advances in graph theory analysis have lead to findings that challenge the notion that functional segregation between regions within default-mode and control networks invariably support cognitive task performance (Cocchi et al., 2013b; Fornito et al., 2012; Hearne et al., 2015). Fornito and colleagues (2012) identified a division of the DMN into core and transitional subsystems where the latter facilitates integration between the core DMN and FPN during goal-directed recollection. Moreover it has been shown that increased cognitive demand during cognitive reasoning is accompanied by a loss of segregation and a progressive enhancement of connectivity between control and default-mode networks (Hearne et al., 2015). Related to this, Hearne et al., (2017) also found that increases in reasoning complexity were associated with greater connectivity and more variable community assignment of the FPN. While these previous studies have focused on direct relationships between select cognitive systems, our whole-brain findings complement this more recent view of a context-dependent coordination of task-positive and task-negative brain systems to support healthy cognitive function. With all of this in consideration, there is compelling evidence suggesting that cognitive control is implemented through the flexible reconfiguration of the FPN as it pertains to a wide range of cognitive domains (Cocchi et al., 2013a; Cole et al., 2013; Fornito et al., 2012; Hearne et al., 2017; Ray et al., 2017; Sheffield et al., 2015).

## Methodological Considerations

While the current study provides novel insights into the dynamic brain network reorganization that support cognitive control processing, it is important to acknowledge potential methodological limitations. Recent studies have shown that movement during fMRI acquisition causes systematic changes in functional connectivity (Power et al., 2014; Power et al., 2015; Satterthwaite et al., 2012). To reduce this potential source of error, we took the precaution of performing a ‘beta-scrubbing’ procedure analogous to scrubbing procedures used in resting-state fMRI studies where beta-images containing frame-wise displacement values greater than 0.5mm motion were excluded from beta-series. Notably, preliminary modularity analyses performed on these data prior to our “beta-scrubbing” procedure aimed at eliminating trials with excess motion were highly consistent with modular partitions currently presented.

The utility of the Power atlas has been well supported within the neuroimaging community, as numerous studies have utilized this set of ROIs for various network connectivity analyses. However, it is important to note that the cognitive labels assigned to their networks that have been subsequently adopted in the current study as ‘cognitive systems’ are based upon reverse inference. Furthermore, the modularity approach employed, Louvain Modularity, relies on the assumption that nodes may only be assigned to a single module. While this is common in modularity algorithms applied to brain imaging data, we recognize the plausibility that multi-functional nodes may be involved in more than one module.

Stringent thresholds applied in the current analysis ensure that only strong, positive connections examined (Zalesky et al., 2016). This is standard practice in the overwhelming majority of graph theoretical analyses, however it should be noted that this step might inadvertently discard neurobiologically relevant information. Furthermore, to support the reliability of the presented results, replication of these analyses using a 10% threshold on functional connectivity graphs was highly consistent. While all modularity values were lower at the 10% threshold, the pattern of results was consistent where task partitions were significantly greater than those of their null and Power partitions. Mutual Information comparing the similarity of partitions from the high and low cognitive control trials of each task also showed a similar pattern of results where the DPX and RiSE Recognition tasks showed significantly greater mutual information than the RiSE Encoding task. Finally, the module composition of brain graphs remained highly consistent between the two thresholds examined, yielding a high mean mutual information score (MI=0.84) and low mean variation of information (VI = 0.1136) across the six brain graphs (representing the two trial types from each of the 3 tasks).

## Conclusions

These findings provide insight into how brain networks reorganize to support cognitive performance under differing task contexts. Results suggest that the FPN can contribute to task appropriate responses through two different mechanisms. Enhanced within-network connectivity in the FPN network is sufficient to support proactive cognitive control, as seen during the DPX. Enhanced network connectivity has also been reported with FPN in the RiSE (Ray et al., 2017), however the FPN also shows a unique capacity for integrating with elements of the DMN and memory network to support different forms of encoding and retrieval.

## Acknowledgments

The authors would like to thank the participants in this study, who gave generously of their time. This research was supported by a research grant from the National Institute of Mental Health (5R01MH059883). All authors approved the final version of the paper for submission.

